# Conformational selection drives substrate-ATP coupling in the type VI ABC transporter

**DOI:** 10.64898/2026.08.11.744245

**Authors:** Samuel Seidl, Aaron Klausnitzer, Jagdeep Kaur, Pelin Zengin, Clemens Glaubitz

**Affiliations:** Institute for Biophysical Chemistry and Center for Biomolecular Magnetic Resonance (BMRZ) Goethe University Frankfurt Max von Laue Straße 9, 60438 Frankfurt am Main, Germany

## Abstract

The mechanism of substrate–ATP coupling is central to the function of ABC transporters. Substrates bind to the specialized transmembrane domains (TMDs), whereas ATP binding and hydrolysis occur in the conserved intracellular nucleotide-binding domains (NBDs). Efficient substrate transport requires coupling between these processes, in which the intracellular coupling helices play an essential role. Here, we probed both coupling helices of the type VI ABC transporter LptB_2_FG(C), the core complex responsible for intermembrane lipopolysaccharide (LPS) transport in Gram-negative bacteria, using site-specific ^19^F labeling and ultra-fast MAS NMR. We show that both coupling helices exist in an equilibrium among three major conformational states. Progression through the coupling cycle occurs via conformational selection, as LPS and nucleotide binding shift the conformational equilibrium, providing evidence for bidirectional communication between the TMDs and NBDs. Furthermore, the conformational distributions of the two coupling helices are asymmetric. Finally, complex formation with LptC, a unique feature of type VI ABC transporters, symmetrizes the conformational landscape of the coupling helices and facilitates transitions between conformational states, thereby enhancing the efficiency of coupling ATP hydrolysis to LPS transport. Together, our findings establish the coupling helices as dynamic allosteric elements that integrate nucleotide and substrate binding through conformational selection, providing a mechanistic framework for substrate–ATP coupling in ABC transporters.

## 1 Introduction

The ATP-binding cassette (ABC) transporter superfamily mediates substrate translocation across biological membranes in all domains of life. Beyond their fundamental physiological functions, ABC transporters are major contributors to multidrug resistance in cancer and antimicrobial resistance in bacteria, making them important therapeutic targets. Over the past decades, extensive structural and biochemical studies have generated a wealth of high-resolution snapshots that capture distinct conformational states along the transport cycle, providing unprecedented insight into their molecular mechanism.^1–3^ ABC transporters share a conserved modular architecture comprising two substrate-specific transmembrane domains (TMDs) and two highly conserved intracellular nucleotide-binding domains (NBDs). Based on their specific topology, 7 classes can be distinguished.^2^ Chemical energy derived from ATP binding and hydrolysis is converted into conformational changes, most notably dimerization of the nucleotide-binding domains (NBDs). Conversely, substrate binding to the transmembrane domains (TMDs) has also been shown to promote NBD dimerization, highlighting the bidirectional allosteric coupling between the two domain types. Several structural elements have been identified as key mediators of communication between the NBDs and TMDs: Within the TMD, one or two coupling helices (CH), depending on the ABC transporter type, establish contacts with the NBDs.^2, 4^ On the NBD side, the Q loop, X loop and the recently identified hinge region mediate interactions with the TMDs.^5^ However, the molecular mechanism underlying the bidirectional crosstalk between NBDs and TMDs remain elusive. In particular, it is unclear how NBD dimerization drives conformational rearrangements in the TMDs and how substrate binding within the TMDs is communicated back to the NBDs to regulate ATPase activity.

Here, we explore the conformational landscape of the coupling helices in the lipopolysaccharide (LPS) transporter LptB_2_FG, representing the only type VI ABC transporter.^2^ LPS is an essential and protective part of the outer membrane of Gram-negative bacteria.^6, 7^ Two LptB proteins constitute the NBD which form a stable complex with the heterodimeric TMD consisting of LptF and LptG.^8^ Apart from TMD-NBD contacts mediated by the CHs, an additional arginine in the intracellular loop 2 of both LptF and LptG form essential contacts to bound ATP.^9^ LPS is extruded by LptB_2_FG and handed over to the single-pass transmembrane protein LptC.^10^ The transmembrane helix (TMH) of LptC intercalates between the TMDs of LptF and LptG, which is a unique feature of the ABC transporter. This is accompanied by an inhibition of ATPase activity of LptB_2_FG^10–12^ and implies an additional regulatory role for LptC.

After transfer to the periplasmic domain of LptC, LPS is pushed to LptA whose subunits form a bridge across the periplasm.^13^ This transport is powered by LptB_2_FG which extrudes LPS downstream^14^ towards LptDE which integrates LPS into the outer leaflet of the outer membrane.^15, 16^ All components, which participate in bridge formation and LptG feature a common β-jellyroll motif for LPS binding.^17^ Due to its essential role in LPS transport, LptB_2_FG is a potential antimicrobial target.^18–21^ Currently, several structures of LptB_2_FG in complex with LPS or in nucleotide-bound states display large structural rearrangements^9, 22, 23^ (Figure 1) but these changes are not reflected in the surroundings of the coupling helices (Figure 1; insets). Therefore, elucidating the mechanism of energy transmission in LptB_2_FGC requires resolving the conformational landscape of the coupling helices.

**Figure 1:**
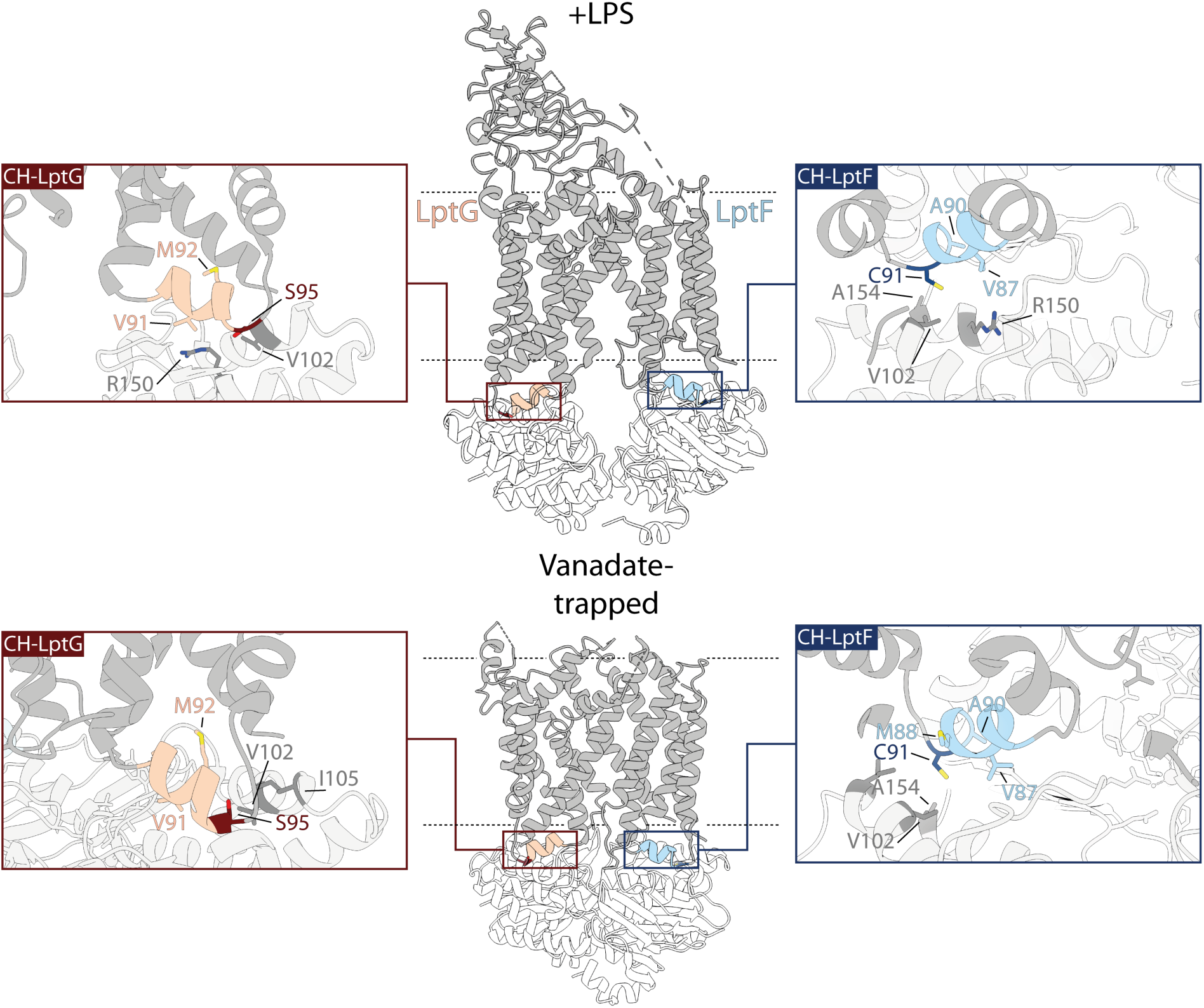
Structures and ^19^F-labelling positions of LptB_2_FG. Previously determined cryo-EM structures of *E. coli* LptB_2_FG co-purified with LPS (top, PDB ID 6MHU ^22^), trapped by ADP-VO_4_ (bottom, PDB ID 6MHZ ^22^). The extracellular β-jr domain was not resolved in the vanadate-trapped structure. The coupling helices of LptG (CH-LptG) and LptF (CH-LptF) are colored in light red and light blue, respectively. The insets show their local environments. LptF-C91 (dark blue) and LptG-S95C (dark red) are the ^19^F TET labelling sites. Surrounding amino acids within a 5 Å radius are depicted. Residues stemming from LptB are colored in dark grey.

The mechanisms how coupling helices work are underexplored. Structural snapshots of ABC transporters only allow a model where the conformational changes of the NBD are rigidly transmitted to the TMD *via* the coupling helices.^24^ Additionally, the coupling helices are shielded constantly by the protein despite the high-amplitude movements of the transporter causing labelling difficulties for large chemical probes e.g. fluorophores or electron paramagnetic (EPR) spin labels. In the worst case, these probes disrupt the sensitive TMD-NBD interface. So far, some studies about coupling helices relied on mutagenesis and biochemical analyses of transport and ATP hydrolysis^25, 26^ or employed atomistic but short MD simulations illuminating initial changes of the coupling helices.^26, 27^ The first direct investigation of the coupling helices using site-specific isotopic amino acid labeling for solid-state NMR demonstrated bidirectional communication in the type IV ABC transporter MsbA.^28^

Here, we asked whether substrate or nucleotide binding shifts the coupling helices between distinct conformations or modulates an underlying conformational ensemble. Given the heterodimeric TMD, we also investigated conformational asymmetry between the coupling helices that could result in differential coupling of ATP hydrolysis to transport. To resolve their conformational landscape, we employed ^19^F ultrafast MAS NMR. The high sensitivity of ^19^F to its local environment enables detection of low-populated conformational states, while MAS at 100 kHz provides high sensitivity and resolution in near-native proteoliposomes.^29, 30^

## 2 Results

### 2.1 The asymmetric conformational landscape of ATP coupling

For ^19^F labelling, two sites were selected: In LptF, based on a cysteine-free construct,^31^ the native cysteine LptF-C91 was used, while in LptG a mutation was introduced (LptG-S95C) (see Figure 1). Both sites were individually labelled with trifluoroethanthiol (TET). Consequently, two samples were prepared: LptB_2_FG(C91^TET^)G and LptB_2_FG(S95C^TET^) to which we refer to CH-LptF and CH-LptG, respectively. The mutants were purified and functionality in terms of ATPase activity, LPS binding and complex formation with LptC was thoroughly assessed (Figure S1). The labelled constructs were then reconstituted into POPE/POPG liposomes and sedimented into the MAS rotor for ^19^F NMR measurements. In order to probe conformational changes within the coupling helices during cross talk between NBDs (LptB) and TMDs (LptF/G), the transporter was prepared in the presence of its substrate LPS, trapped in the pre-hydrolysis state by AMP-PNP or both. Thus, distinct states during coupling could be characterized.

The fluorine spectra of CH-LptF and CH-LptG in the apo state are both characterized by a complex line shape stretching from approx. 7 ppm to 12 ppm (Figure 2). Spectral deconvolution reveals three distinct peaks, which may three different coupling helix-related protein conformations. The results indicate that the coupling helices sample a highly diverse structural environment.

**Figure 2:**
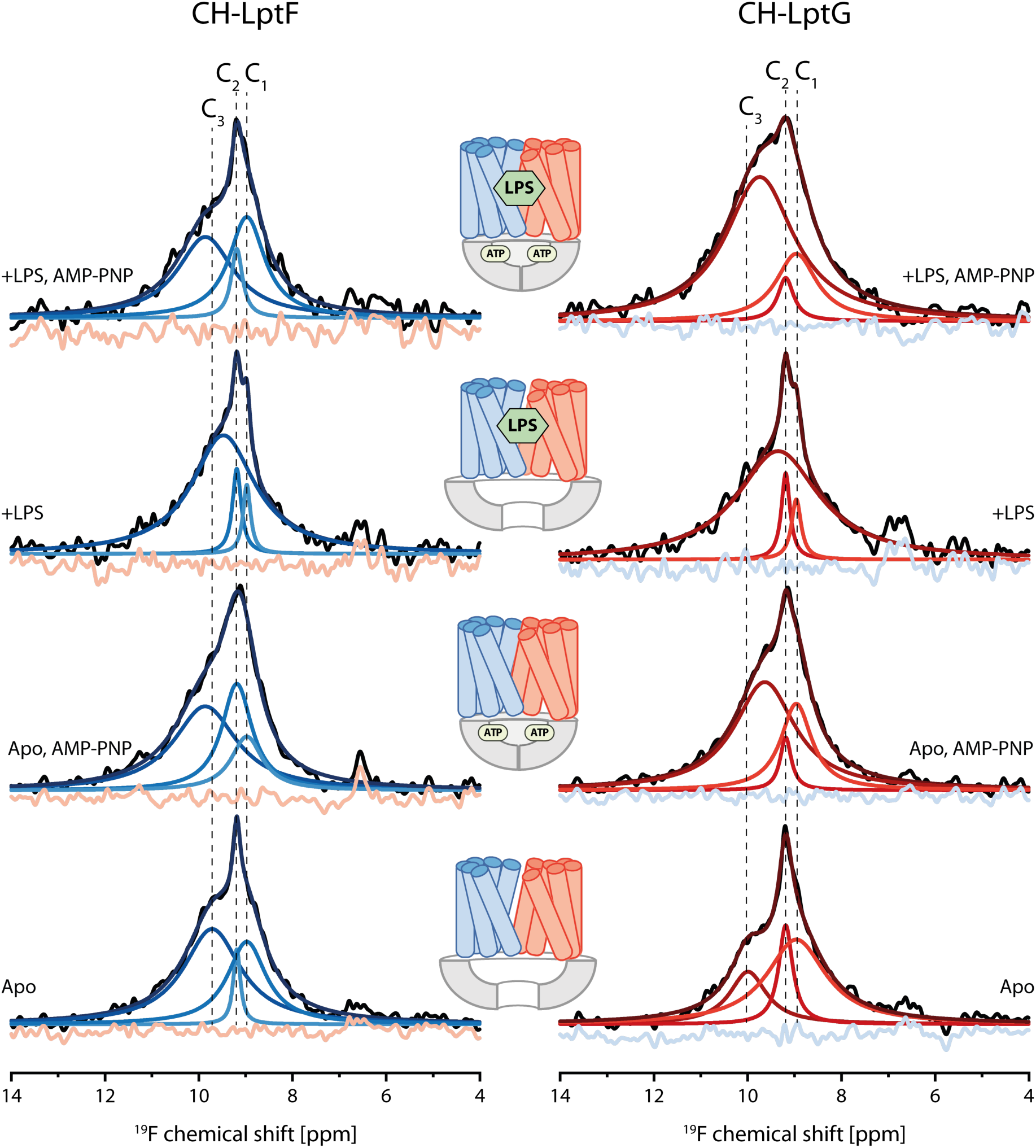
^19^F-MAS NMR spectra of labelled LptB_2_FG represent the conformational landscape of both coupling helices in LptF and LptG. **(A)** 1D ^19^F spectra of LptB_2_F(C91^TET^)G (CH-LptF) in the apo state without LPS (bottom), without LPS but bound to AMP-PNP (bottom middle), co-reconstituted with LPS (top middle) and co-reconstituted with LPS and bound to AMP-PNP (top). **(B)** As in (A) but for LptB_2_FG(S95C^TET^) (CH-LptG). The experimentally acquired spectra are shown in black. Deconvolution was performed by fitting three Lorentzian functions to the experimental line shapes. The individual deconvoluted peaks are shown in lighter colors, while the overall fit is displayed in dark blue or dark red, respectively. The residuals are depicted in light blue or light red, respectively. Deconvolution results are summarized in Tables S1, S2 and Figure S2.

Although the TMD is heterodimeric, the lineshapes and chemical shifts observed for CH-LptF and CH-LptG are highly similar, indicating that both coupling helices experience comparable chemical environments. These findings suggest that the two TMDs sample the same major conformational states but differ in the population distributions of these states at their respective coupling helices. Because only a single probe is monitored in each spectrum, the presence of multiple resolved peaks indicates that exchange between the conformations is slow on the NMR timescale.

For CH-LptF (Figure 2A), spectral deconvolution yields chemical shift differences of 0.53 ppm between C_3_ and C_2_ and 0.22 ppm between C_2_ and C_1_, corresponding to frequency separations of 424 Hz and 176 Hz, respectively. These values indicate that exchange between the corresponding conformational states occurs on timescales slower than approximately 2 ms and 5 ms, respectively. Furthermore, the three deconvoluted peaks exhibit distinct linewidths and relative intensities. The most populated state is C_3_ (59 %) followed by C_1_ (35%) and C_2_ (6%). Peak linewidths can provide additional information about the dynamics of the corresponding conformational state, the presence of unresolved substates, or both. Here, C_3_ displays the broadest linewidth (1.48 ppm), followed by C_1_ (1.00 ppm) and C_2_ (0.18 ppm) (Table S1).

Although the apo spectrum of CH-LptG (Figure 2B) initially appears similar to that of CH-LptF (Figure 2A), spectral deconvolution reveals notable differences. Peak C_3_ is shifted by −0.27 ppm relative to CH-LptF, whereas the remaining peaks exhibit nearly identical chemical shifts (Table S2; Figure S2A). This small chemical shift difference implies a shift of unresolved substates within C_3_ between CH-LptG and CH-LptF. In contrast to CH-LptF, C_1_ represents the predominant population (60%), followed by C_3_ (23%) and C_2_ (17%) (Table S2). The linewidths also differ from those observed for CH-LptF: C_3_ is narrower, whereas C_2_ and C_1_ become broader.

Having established that three conformations are already populated in the apo state, we next characterized how each conformation responds to the addition of either the substrate LPS or the ATP analogue AMP-PNP, which traps the transporter in the pre-hydrolysis state or both.

Trapping the transporter in the pre-hydrolysis state with AMP-PNP results in broader spectra for both samples (Figure 2). For CH-LptF, C_3_ undergoes a shift of +0.14 ppm relative to the apo state (Figure 2A). In addition, the populations of the three conformations are redistributed. Most notably, the population of C_2_ increases from 6% to 31%, whereas that of C_1_ decreases from 35% to 16%. The linewidths of both C_2_ and C_3_ increase, while that of C_1_ decreases. Interestingly, CH-LptG exhibits a distinct response to AMP-PNP trapping (Figure 2B). In contrast to CH-LptF, C_3_ undergoes a shift of -0.35 ppm relative to the apo state. Moreover, the population of C_3_ increases from 23% to 67%, whereas the populations of C_2_ and C_1_ decrease from 17% to 6% and from 60% to 27%, respectively. This goes along with a larger linewidth for C_3_ but narrower peaks for C_1_ and C_2_.

We next investigated how substrate binding influences the conformational dynamics of the coupling helices. To this end, LPS was co-reconstituted with LptB_2_FG to probe substrate-dependent effects (Figure 2). Under these conditions, the spectra of CH-LptF and CH-LptG become more similar. In both cases, the population of C_3_ increases further, reaching 86% and 87% for CH-LptF and CH-LptG, respectively. C_1_ and C_2_ populations decrease and their peaks become narrower, exhibiting very similar intensities and linewidths. Relative to the apo state, C_3_ undergoes a shift of -0.25 ppm in CH-LptF and -0.64 ppm in CH-LptG.

To mimic the pre-hydrolysis state, in which the nucleotide-binding domains (NBDs) dimerize and the LPS-binding pocket collapses to drive LPS extrusion, we combined LPS binding with AMP-PNP trapping (Figure 2). Under these conditions, the spectra of CH-LptF and CH-LptG diverge markedly. The relative populations of C_3_, C_2_, and C_1_ are 51%, 9%, and 40% for CH-LptF, compared with 74%, 5%, and 21% for CH-LptG. The CH-LptF spectrum closely resembles that of the apo state with respect to the conformational populations, whereas the chemical shift of C_3_ remains similar to that observed in the AMP-PNP-trapped state. In contrast, the CH-LptG spectrum closely resembles the AMP-PNP-trapped state in terms of conformational populations, linewidths, and chemical shifts. The only notable difference is a shift of the C_3_ resonance by +0.10 ppm. Relative to the LPS-bound state, however, the C_3_ resonance is shifted by +0.39 ppm in both CH-LptF and CH-LptG.

Interestingly, these data show that the similar conformational equilibria of the coupling helices in LptF and LptG observed in the LPS-bound state become highly asymmetric upon further nucleotide binding.

### 2.2 The influence of LptC on ATP coupling

In order to investigate the influence of LptC on the coupling helices, the same sample conditions as before are applied, but the full LptB_2_FGC complex was *in vitro* assembled as previously described ^11^ and ^19^F spectra of TET-labelled coupling helices were recorded. We refer to these samples LptB_2_F(C91-TET)GC and LptB_2_FG(S95C-TET)GC in the following as CH-LptF+C and CH-LptG+C, respectively.

The apo spectra of CH-LptF+C and CH-LptG+C differ substantially from those recorded in the absence of LptC and can be well described by deconvolution into three Lorentzian peaks, C_3_, C_2_, and C_1_, as outlined above (Figure 3; Tables S3 and S4). In contrast to the apo spectra of CH-LptF and CH-LptG, the spectra of CH-LptF+C and CH-LptG+C are highly similar, exhibiting almost identical chemical shifts and comparable conformational populations, with C_3_: C_2_: C_1_ ratios of 48:8:43 and 52:8:41, respectively. Furthermore, all three peaks display increased linewidths in the presence of LptC. Compared with CH-LptF, CH-LptF+C exhibits small chemical shift perturbations for C_3_ and C_2_, whereas in CH-LptG+C only C_3_ is slightly shifted relative to CH-LptG (Tables S1–S4).

**Figure 3:**
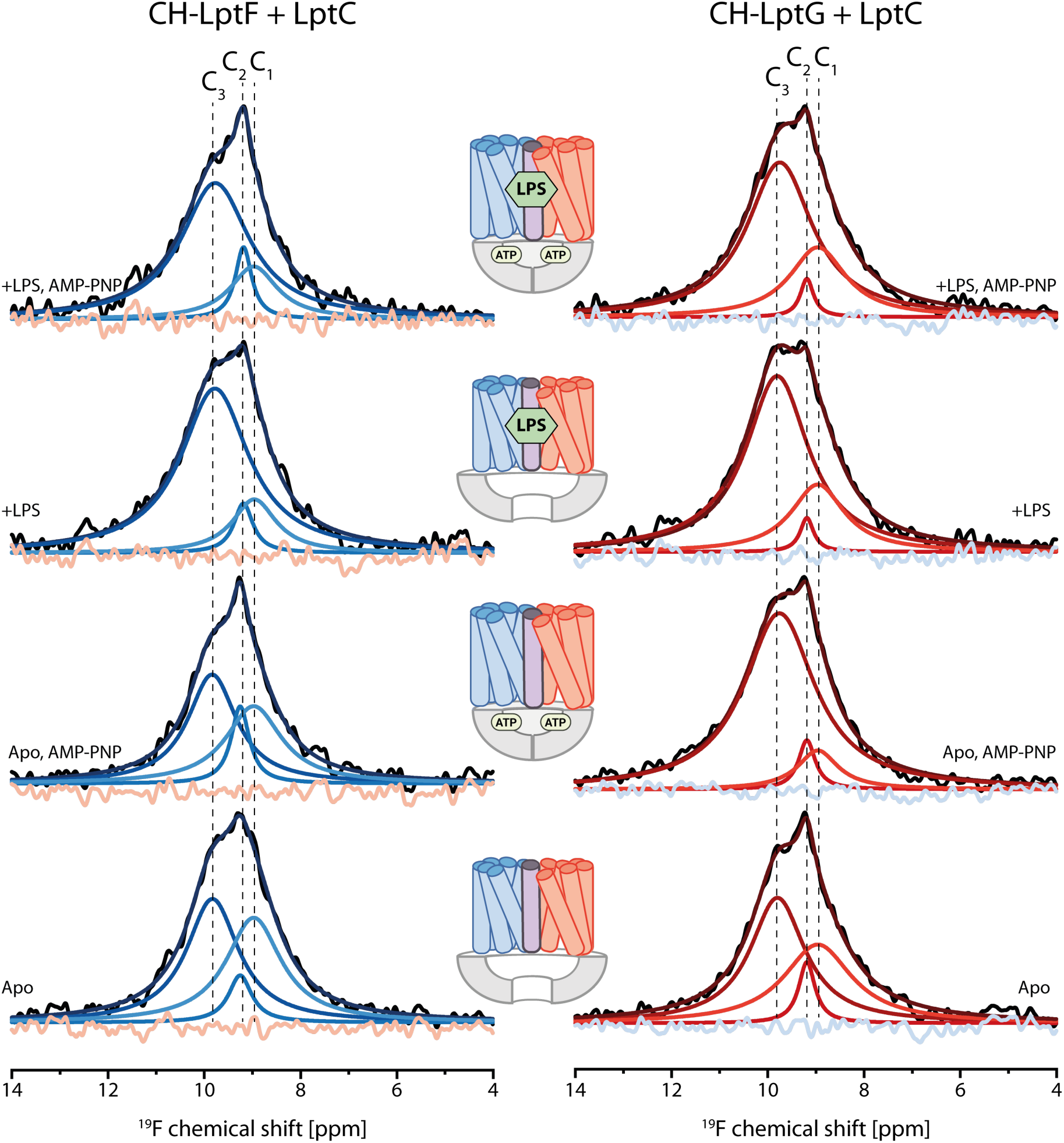
Effect of LptC on the ^19^F-MAS NMR spectra and the conformational landscape of both coupling helices in LptF and LptG in LptB_2_FG. **(A)** 1D ^19^F spectra LptB_2_F(C91^TET^)GC (CH-LptF+C) in the apo state without LPS (bottom), without LPS but bound to AMP-PNP (bottom middle), co-reconstituted with LPS (top middle) and co-reconstituted with LPS and bound to AMP-PNP (top). **(B)** As in (A) but for LptB_2_FG(S95C^TET^)C (CH-LptG+C). The experimentally acquired spectra are shown in black. Deconvolution was performed by fitting three Lorentzian functions to the experimental line shapes. The individual deconvoluted peaks are shown in lighter colors, while the overall fit is displayed in dark blue or dark red, respectively. The residuals are depicted in light blue or light red, respectively. Deconvolution results are summarized in Tables S3, S4 and Figure S2.

Next, the apo, AMP-PNP-trapped state was examined. Under these conditions, the spectra of CH-LptF+C and CH-LptG+C differ substantially, most notably in the relative populations of C_3_, C_2_, and C_1_ (48:13:39 vs. 84:6:11, respectively). These data indicate that, in the presence of LptC, the coupling helix of LptG responds more strongly to AMP-PNP binding than that of LptF, with the conformational equilibrium shifting predominantly toward the C_3_ state. Compared with the corresponding sample lacking LptC, the spectrum of CH-LptF+C remains largely unchanged, whereas CH-LptG+C exhibits a pronounced redistribution of the conformational ensemble.

In the LPS-bound state, the spectra of CH-LptF+C and CH-LptG+C again become highly similar, exhibiting comparable conformational populations with C_3_: C_2_: C_1_ ratios of 77:7:16 and 73:3:24, respectively. A similar trend was observed in the absence of LptC; however, the C_3_ state was even more highly populated, while the C_1_ and C_2_ populations became smaller.

For the LPS-bound, AMP-PNP-trapped transporter, the chemical shifts of the C_3_ resonances in CH-LptF+C and CH-LptG+C remained similar to those observed in the LPS-bound state. Interestingly, the conformational populations also closely resembled those of the LPS-bound, nucleotide-free transporter. In CH-LptF+C, the population of C_3_ decreased slightly to 72%, accompanied by a modest increase in the C_2_ and C_1_ populations to 9% and 19%, respectively. For CH-LptG+C, the conformational populations were essentially identical to those of the LPS-bound, nucleotide-free state, mirroring the behavior of CH-LptF+C. In contrast to the transporter lacking LptC, however, the conformational populations of the three states were considerably more symmetric between the LptF and LptG coupling helices in the presence of LptC.

## 3 Discussion

Using site-specific ^19^F labeling and ultra-fast MAS NMR, we found that both CHs populate three distinct conformational states whose relative populations are redistributed upon nucleotide binding, LPS binding, combined LPS and nucleotide binding, and in the presence of LptC. All observed spectroscopic changes are summarized in Figure S2.

### The coupling helices undergo conformational exchange in the apo state

Based on the available structural snapshots of ABC transporters, conformational changes in the NBDs appear to be transmitted rigidly to the TMDs through the coupling helices and a conserved arginine residue in intracellular loop 2 of LptF or LptG.^9, 32^ In such a model, the coupling helices would adopt a single conformation in each functional state, resulting in a single, well-defined resonance that shifts upon transitions between states. Instead, our data reveal a markedly different picture. In the apo state, the spectra of both coupling helices exhibit complex lineshapes that can be deconvoluted into multiple overlapping resonances. Because each sample contains only a single ^19^F label, the presence of multiple resonances demonstrates the coexistence of distinct conformational states that exchange slowly on the NMR timescale. Based on the measured *v*_max_ values of the labeled mutants, the transporter hydrolyzes approximately one ATP molecule every three seconds (Figure S1B). Thus, the slow conformational exchange observed by NMR is fully compatible with the catalytic turnover rate.

The similar lineshapes and chemical shift ranges of CH-LptF and CH-LptG indicate that both coupling helices sample the same major conformational states. However, differences in linewidths, populations, and C_3_ chemical shifts in the apo state reveal distinct conformational equilibria for CH-LptF and CH-LptG.

### The coupling process is asymmetric and based on conformational selection

Based on the complete set of spectra, we propose a model in which the coupling helices populate three distinct conformational states. Each of these states is likely associated with a specific stage of the ATPase/transport cycle and thus represents a discrete step in the coupling of NBD motions to the TMDs. The distinct responses of the three peaks to the different experimentally prepared states (Figure S2) enable their assignment to specific stages of the ATPase/transport cycle.

The C_3_ state is enriched for both coupling helices in the presence of LPS or AMP-PNP. Its enrichment in the AMP-PNP-bound state allows C_3_ to be assigned to the coupling-helix conformation of the AMP-PNP-bound structure.^9^ The C_3_ peak changes not only in integral intensity but also in chemical shift and linewidth upon binding of either LPS or AMP-PNP. The observed chemical shift changes indicate that C_3_ samples multiple unresolved substates, whose populations differ between these binding conditions. Consequently, LPS binding shifts the conformational ensemble toward a different substate than that populated in the presence of AMP-PNP alone.

In the case of LPS, these observations may indicate feedback coupling mediated by the coupling helices, which stabilizes a conformational substate in which the NBDs are positioned closer together prior to nucleotide-induced dimerization. This pre-organized state could promote ATP hydrolysis, consistent with mechanisms proposed for LptB₂FG and other ABC transporters.^11, 33–37^ However, currently available LPS-bound and apo structures of LptB₂FG do not reveal clear differences in coupling helix orientations.^22, 38^ It remains unclear whether current structural methods are capable of resolving these proposed transporter substates.

The observed increase in linewidth may indicate the presence of additional substates, consistent with the substrate-dependent equilibrium chemical shifts observed for C_3_. However, the observed linewidth reflects a complex interplay between the presence of multiple conformational substates and dynamic processes on the intermediate exchange timescale. T₂ relaxation experiments are therefore required to disentangle these contributions.

Peak C_2_ exhibits only minor changes in population and linewidth across the different conditions (Figure S2). The only notable deviation is observed in LptF in the AMP-PNP-trapped state, although no consistent trend can be identified. The limited response of C_2_ to substrate or nucleotide binding suggests that it represents an intermediate conformation between the apo and nucleotide-trapped states. Such an intermediate could serve as a gating state for successful coupling, with productive coupling requiring sequential passage through all three conformational states before returning to the initial state.

Peak C_1_ exhibits behavior opposite to that of C_3_ and therefore likely represents an apo-like conformation with separated NBDs. LPS binding is accompanied by an enrichment of C_3_ and a marked depletion of C_1_, suggesting that the conformational equilibrium shifts away from the apo-like state. One possible interpretation is that LPS binding results in a thermodynamically unfavorable situation which is stabilized by depletion of C_1_. Thereby return to the apo state is blocked which leads to a halt of the coupling cycle. Such a mechanism could prevent uncoupled LPS extrusion in the absence of ATP hydrolysis.

As a result, it can be deduced that prior to the transport against a gradient, the extrusion is the energy-demanding step. This ATP-dependent ejection of substrate was recently demonstrated for another type IV ABC transporter.^39^ Here, we could show that the suppression of CH orientations is the underlying cause. Taking both observations into account, this could be a principle which concerns all ABC transporters.

As all three conformations are present under every experimental condition, our data suggest that the coupling helices move largely independently of the nucleotide-binding domains (NBDs). Consequently, ATP binding therefore does not induce the conformational transition itself; instead, it biases the pre-existing equilibrium toward a specific coupling helix conformation, supporting a conformational selection mechanism for ATP coupling (Figure 4A). It also means that substrate–ATP coupling does not achieve 100% efficiency. This model may also explain why the transport cycle can require the hydrolysis of more than two ATP molecules to translocate a single substrate molecule.^40–42^

**Figure 4:**
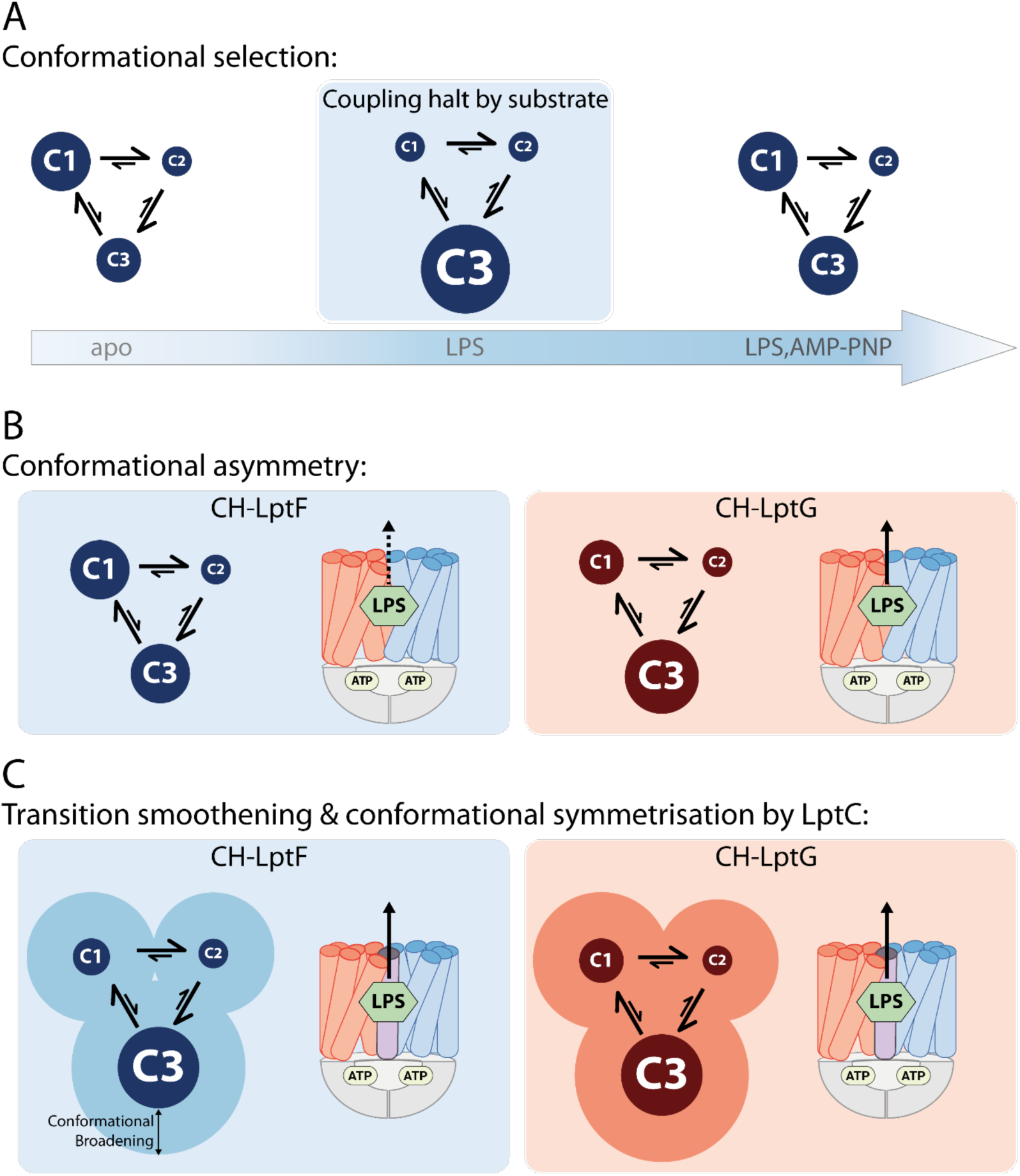
Summary of the shift of the conformational equilibria of both coupling helices by the binding of LPS and nucleotide. **(A)** Conformational selection enables progression through the coupling cycle: In the apo state, a dynamic equilibrium among three conformational states C_1_, C_2_, and C_3_ exists. Upon substrate binding, the equilibrium shifts toward C_3_. Subsequent nucleotide binding resets the equilibrium and enabling continued progression through the coupling cycle. The representative scheme shown here illustrates the conformational populations of the LptF coupling helix in LptB_2_FG. The size of each circle is proportional to the population of the corresponding conformational state (see Figure 2A and Table S1). **(B)** The conformational equilibria of the LptF and LptG coupling helices are asymmetric within LptB_2_FG (see Figures 2A, B and Tables S1, S2). The LptG coupling helix preferentially populates the C_3_ state, whereas the LptF coupling helix exhibits a more balanced distribution between the C_1_ and C_3_ states. Consequently, ATP binding is expected to more effectively promote LPS transport through the LptG subunit. **(C)** The presence of LptC symmetrizes the conformational equilibria of the LptF and LptG coupling helices (Figures 3A, 3B; Tables S3, S4), resulting in more similar conformational populations between the two subunits. This increased symmetry suggests that the coupling helices operate in a more coordinated manner, thereby promoting more efficient coupling of ATP hydrolysis to LPS transport.

The changes observed in both coupling helices are significantly asymmetric which appears plausible for an ABC transporter with a heterodimeric TMD. C_3_ is most of the time more enriched in LptG than in LptF especially in the nucleotide-trapped state. Moreover, C_1_ is more populated in CH-LptF. Consequently, CH-LptG seems to progress more efficiently through the coupling cycle. Prior studies hypothesized that LptF plays a more dominant role in the coupling process.^32^ However, we show that in a transporter devout of LptC, CH-LptG promotes LPS extrusion (Figure 4B).

Although the coupling helices contribute essentially to ATP coupling, it is not the sole contributor. The role of the single arginine in loop 2 is undetermined. They form essential contacts to the nucleotide and were determined to be necessary for LPS transport. Consequently, they might also play a role in the coupling process.^43^

### LptC enhances the conformational landscape and smoothens transitions

LptC is required for the transfer of LPS from LptB_2_FG to LptA.^10, 44^ In addition, LptC downregulates the ATPase activity of the transporter. This suggests that LptC may exert allosteric control over/via the coupling helices and thereby influence their conformational dynamics. To test this possibility, the effect of LptC on the conformational landscape of the coupling helices was investigated.

Most strikingly, in the presence of LptC, the coupling helices of LptF and LptG exhibit similar populations of the three conformational states (C_1_– C_3_) relative to each other across all experimentally prepared sample conditions. This means that LptC symmetrizes the sampling of the conformational states of both coupling helices and that LptF works in a synchronized fashion together with LptG. This is in contrast to the situation without LptC where LptG seems to play a more dominant role as discussed above. Our findings suggest that LptC enhances the coupling efficiency between ATP hydrolysis and LPS transport making LPS extrusion more efficient (Figure 4C).

Interestingly, in the LPS-bound state, the linewidths of the C_1_ and C_2_ peaks, corresponding to the apo-like and intermediate conformational states, broaden markedly in the presence of LptC compared with the corresponding state in its absence, indicating enhanced conformational dynamics, which is compatible with previous reports.^9, 12^ This increase in dynamics may facilitate transitions between the major conformational states. Together with the observed symmetry of conformational sampling, these findings suggest that LptC promotes a more efficient coupling between ATP hydrolysis and LPS transport.

## 4 Summary

We showed that both coupling helices exist in an equilibrium among three major conformational states under all experimentally prepared conditions, indicating that substrate–ATP coupling in the type VI ABC transporter is a highly dynamic process. ATP binding does not dictate a specific conformational transition but instead shifts a pre-existing conformational equilibrium, enabling the coupling helices to progress through the coupling cycle through **conformational selection**. This mechanism provides a plausible explanation for the intrinsic inefficiency of substrate–ATP coupling. LPS further modulates the equilibrium by promoting progression through the transport cycle, providing evidence for bidirectional communication mediated by the coupling helices. Such a conformational selection mechanism may also operate in other ABC transporters, particularly those that undergo “futile” ATP hydrolysis cycles.

Furthermore, the conformational distributions of the LptF and LptG coupling helices is **asymmetric**, indicating that the two subunits transmit conformational changes from the nucleotide-binding domains (NBDs) to the transmembrane domains (TMDs) with different efficiencies. In the context of LptB_2_FG, this asymmetry is particularly intriguing in light of LptB_2_FG mutants that bypass the requirement for LptC during LPS transfer.^45^ In other heterodimeric ABC transporters, however, such asymmetric coupling may have even more direct functional consequences.

Finally, complex formation with LptC - a unique feature of type VI ABC transporters - **symmetrizes** the conformational landscape of the coupling helices and facilitates transitions between conformational states, thereby enhancing the efficiency of coupling between ATP hydrolysis and LPS transport.

## 5 Limitations of this study

The observations we made here describe the coupling process in type VI ABC transporters. We cannot rule out that the mechanism differs in other ABC transporter types. Furthermore, the allosteric effects from downstream components of the Lpt system were not addressed.^44, 46^

## 6 Acknowledgements

This work was funded by the CRC 1507 ‘Membrane-associated protein assemblies, machineries and supercomplexes’ and by the LOEWE project GLUE (G protein-coupled receptor Ligands for Underexplored Epitopes) of the State of Hesse. We thank Anouk Ebenzer, Goethe University Frankfurt for help with sample preparations and Dr. Melanie MacDowell, MPI for Biophysics Frankfurt, for supporting our NanoDSF experiments.

