## Supporting information and data for "Conformational selection drives substrate-ATP coupling in the type VI ABC transporter"

Goethe University Frankfurt

### Material and Methods

#### Generation of Mutants

Mutants were generated using *E. coli* $LptB_2FG$ in pETDuet-1 (Dajka et al., 2024). Quikchange multi-site directed mutagenesis was used according to the manufacturers protocol with the primer listed in the materials table. After mutagenesis, parental DNA was digested using DpnI and 2.5 µl of the digest was used to transform *E. coli* DH5α cells, streaked on LB-agar plates containing ampicillin and incubated at 37 °C overnight. Then, colonies were selected and incubated in 5 ml LB containing ampicillin at 37 °C overnight. Next, DNA was isolated from these cultures using MN Nucleospin plasmid. The isolated DNA was sent for DNA sequencing to verify the mutation.

#### Expression and Purification

**Expression and purification of $LptB_2FG$.** For transformation of $LptB_2FG$ in *E. coli*, 1 µl of plasmid DNA was mixed with 50 – 100 µl suspension of chemically competent *E. coli* C43 (DE3). The suspension was incubated for 15-30 min on ice. Then, the suspension was incubated at 42 °C for 90 sec. After cooling down on ice for 5 min, the suspension was diluted with LB medium without antibiotics to 1 ml and the suspension was shaken at 37 °C for 1 h at 550 rpm. Subsequently, 100 µl was spread onto LB-agar plates and the plates were incubated at 37 °C overnight. Then a colony was selected to inoculate 100 µ of LB medium containing ampicillin and this pre-culture was shaken at 37 °C at 180 rpm overnight. Then, 10% of the main culture volume was taken to inoculate 500 ml of M9 medium (composition: 3.39 g $K_2HPO_4$, 1.5 g $KH_2PO_4$, 0.25 g NaCl, 0.5 g $NH_4Cl$, 2 g glucose, 1.0 mL of 1 M $MgSO_4$, 0.5 mL of 0.01 M $FeCl_2$ per 500 mL culture and all amino acids) in a 2 l shaking flask. At an $OD_{600}$ of 0.4 temperature was reduced to 18°C. Between 0.6 and 0.8 protein expression was induced by adding 0.2 mM IPTG. Expression was carried out between 16 and 18 hours at 18 °C at 185 rpm.

The cell pellet from 0.5 liter culture was resuspended in 50 ml Lpt lysis buffer containing protease inhibitor and DNAse I and lysed using a high-pressure homogenizer cycling the lysate thrice at a pressure of 1.8 kbar. The lysate was then centrifuged at 5000 × g for 10 min at 4 °C in order to separate larger cell fragments and unbroken cells. The supernatant was then subjected to ultracentrifugation at 250,000 × g for 1 h or 200,000 × g for 1:30 h, both at 4 °C. For solubilization each gram of membrane (if not indicated otherwise), was resuspended in 10 ml Lpt lysis buffer containing protease inhibitor and 1% *n*-dodecyl-β-D-maltoside (DDM) with 2 mM ATP. The suspension was stirred for 1 h at 4 °C. Afterwards, the suspension was ultracentrifuged at 250,000 × g for 1 h at 4 °C. The supernatant was then filtered using a 0.45 µm syringe filter and applied to a HisTrap HP column running at 1 ml/min. After the flow-through, the column was washed with Lpt purification buffer until the absorption at 280 nm ($A_{280}$) was at a baseline level. Then, the column was washed with 20 CV (column volumes) Lpt wash 1 and 15 CV Lpt wash 2. The protein was then eluted with 10 CV Lpt elution buffer.

For $^{19}F$ labelling, 1 mM tris(2-carboxyethyl)phosphine (TCEP) was added during solubilization. For trifluoroethanthiol (TET) labelling, the labelling was optimized for the purification with the HisTrap HP column. After washing, the column was incubated with 5 CV Lpt purification buffer supplemented with 1 mM 4-DPS and the buffer was circulated for 1 h at 4 °C. After activation of the cysteines, the resin was washed with 15 CV Lpt purification buffer and then incubated with 5 CV Lpt purification buffers supplemented with 1 mM TET. The buffer was circulated for 1 h at 4 °C. Then the column was again washed with 15 CV Lpt purification buffer and the protein was eluted.

**Expression and purification of LptC.** If not indicated otherwise, the expression and purification procedure was performed according to the $LptB_2FG$ protocol. Transformation was performed with *E. coli* BL21 (DE3) cells using chloramphenicol for selection. After the initial LB preculture, a second 0.5-liter M9 preculture was inoculated with 10% of the M9 preculture volume and grown for 18 hours at 37°C at 190 rpm. In the M9 media of LptC the 2 g glucose were substituted with 2 g glycerol. For the main culture the temperature was reduced to 20 °C at an $OD_{600}$ of 0.4. Afterwards, the induction was performed with 0.2% arabinose at an $OD_{600}$ 0.6-0.7 and the cells were grown for 18 hours at 20 °C at 190 rpm.

For the solubilization of LptC, 1% of sodium lauroyl sarcosinate (LS) was added to the LptC purification buffer together with protease inhibitor. After solubilization and centrifugation, the supernatant was supplemented with 5 mM imidazole and applied to Ni-NTA beads for a 1 hour incubation at 4°C. Washing was carried out by 20 CV of LptC purification buffer containing 5 mM imidazole, followed by 15 CV LptC purification buffer containing 20 mM imidazole. The protein was eluted with 3-4 CV LptC purification buffer containing 350 mM imidazole.

### Reconstitution into liposomes

1-palmitoyl-2-oleoyl-sn-glycero-3-phosphoethanolamine/1-palmitoyl-2-oleoyl-sn-glycero-3-phospho-(1′-rac-glycerol (POPE/POPG, 4:1 mol/mol) was dissolved in Chloroform/Methanol (9:1) until the solution was clear. The solution was then dried under dry nitrogen until no visible fluid was left. For complete removal of solvent, the lipids were further dried using a rotary evaporator at 40 mbar overnight. The dried lipids were resuspended in the corresponding lipid buffer so that a concentration of 4 mg/ml is reached. In order to resuspend the lipids, the suspension was rotated in water bath which was above the phase transition temperature of the lipids. The suspension of the lipids was supported by a sonication bath. The lipids were then freeze-thawed three times and then passed through a 0.2 µm polycarbonate membrane using an extruder 11 to 13 times.

Prior to reconstitution, the lipid suspension was destabilized with 4 mM DDM incubating the suspension for 5 min at room temperature. Then, purified $LptB_2FG$ with a concentration of 1 mg/ml was mixed with POPE/POPG at a lipid-to-protein ratio of 100:1 (mol/mol) and mixed at room temperature for 30 min. In case of LPS addition, LPS at a protein-to-LPS-ratio of 1:7.5 (mol/mol) was added 2:30 min prior to protein addition. Then, the detergent was removed using BioBeads three times at a concentration of 80 mg/ml of reconstitution mixture (without considering LPS). For the first time, the mixture was incubated at 4 °C for 1 h. After another addition of BioBeads, the suspension was incubated at 4 °C overnight. The next day, a final round of BioBeads was added and the suspension was incubated again at room temperature for 2 h. The suspension was then filtered using a cell strainer to remove the BioBeads. The suspension was further diluted with Lpt lipid buffer and the proteoliposomes were harvested at 250,000 × g for 1 h at 4 °C and resuspended in Lpt NMR buffer. The proteoliposomes were then washed another two times with Lpt NMR buffer at 82,000 × g and 4 °C for 30 min to remove imidazole. For ATPase assays in proteoliposomes, the proteoliposomes were not harvested to maintain a homogenous suspension.

For AMP-PNP trapping, 10 mM AMP-PNP from a 100 mM stock solution (in HEPES pH 7.5) were added to 400 µg of protein in proteoliposomes. The mixture was subjected to three freeze–thaw cycles (liquid nitrogen and 37 °C water bath) to enhance trapping efficiency, followed by incubation at room temperature for 60 min. Following incubation, the sample was pelleted and washed thoroughly with buffer containing 50 mM HEPES and 5 mM MgCl2 to remove excess reagents.

### Determination of ATPase activity

The ATPase assay is based on Morbach et al. (41). 1 µg of proteoliposomes in ATPase buffer was incubated with different concentrations of ATP (in ATPase buffer) for 1 h at 37 °C at 500 rpm. After incubation, the reaction was stopped with 175 µl 20 mM sulfuric acid. Then, 50 µl of malachite green working solution was added and 80 µl of 11% Tween and 7.5% $NH_4MO_7$. The mix was incubated for 8 min at room temperature and then the absorbance at 620 nm was measured.

### Resolubilization of $LptB_2FGC$

The harvested proteoliposome pellet was resuspended to a protein concentration of 2.3 mg/ml with Lpt lysis buffer. After resuspension, 0.5% DDM was added and incubated for 1 h at 4°C under agitation for resolubilization. To remove insolubilized liposomes the solution was centrifuged at 183,960 × g for 1 h. The samples were concentrated to a volume of 500 µl, achieving a concentration of 1 – 2 mg/ml. Subsequently, samples were analyzed on a Superdex 200 Increase 10/300 equilibrated with Lpt purification buffer.

### Determination of thermal stability

The nanoDSF measurements were recorded using a Prometheus Panta (NanoTemper Technologies, Munich, Germany). Prior to the measurements, $LptB_2FG$ mutants were incubated for 30 minutes at room temperature either in the absence or presence of a 2-fold molar excess of Ra-LPS. For nanoDSF, a temperature gradient of 1 °C/min from 25 °C to 80 °C was applied. Intrinsic tryptophan fluorescence was monitored at 330 nm and 350 nm. Data analysis was performed using the manufacturer's software, and thermal transitions were identified by plotting the first derivative of the 350 nm fluorescence intensity curves.

### NMR spectroscopy

All experiments were performed on a Bruker Avance III wide-bore ssNMR spectrometer operating at 850 MHz $^{1}$H Larmor frequency (800 MHz $^{19}$F Larmor frequency) equipped with a Bruker 0.7 mm HCN probehead tunable to $^{19}$F. Experiments were performed at 100 kHz MAS. Spectra were recorded at 290 K set temperature which corresponds to approximately 25 °C determined by the proton chemical shift of water in the sample. Spectra were referenced to 0.5 mM buffered TFA for $^{19}$F experiments or 3 mM DSS for $^{1}$H experiments which was added before sedimentation into the MAS rotor.

$^{19}$F spectra were acquired using a Hahn echo pulse sequence with a inter pulse delay of 100 µs in order to reduce background signals from the probehead. 90° pulse duration was 1.05 µs. Typically, 50,000-200,000 transients for $LptB_2FG$ with acquisition time of 50 ms. During processing, the FID was typically cut after 10 ms and 75 Hz exponential linebroadening was applied.

**Deconvolution.** Spectra were deconvoluted using Origin Pro 2025 by applying Lorentzian peak fitting. The minimum number of peaks required to adequately reproduce the experimental spectrum was used for the deconvolution. For the apo spectra, three peaks were sufficient to achieve a satisfactory fit, indicating the presence of three major conformational states of the coupling helices, designated $C_1$, $C_2$, and $C_3$. To ensure comparability across all spectra, the chemical shifts of $C_1$ and $C_2$ were determined from the LPS-containing samples, where these peaks were best resolved. These chemical shifts were then held constant during the deconvolution of all spectra, provided that the experimental spectra could be adequately reproduced. In contrast, the chemical shift of the broader $C_3$ peak was allowed to vary during the fitting procedure to account for potential shifts arising from changes in the distribution or preference of its underlying conformational substates.

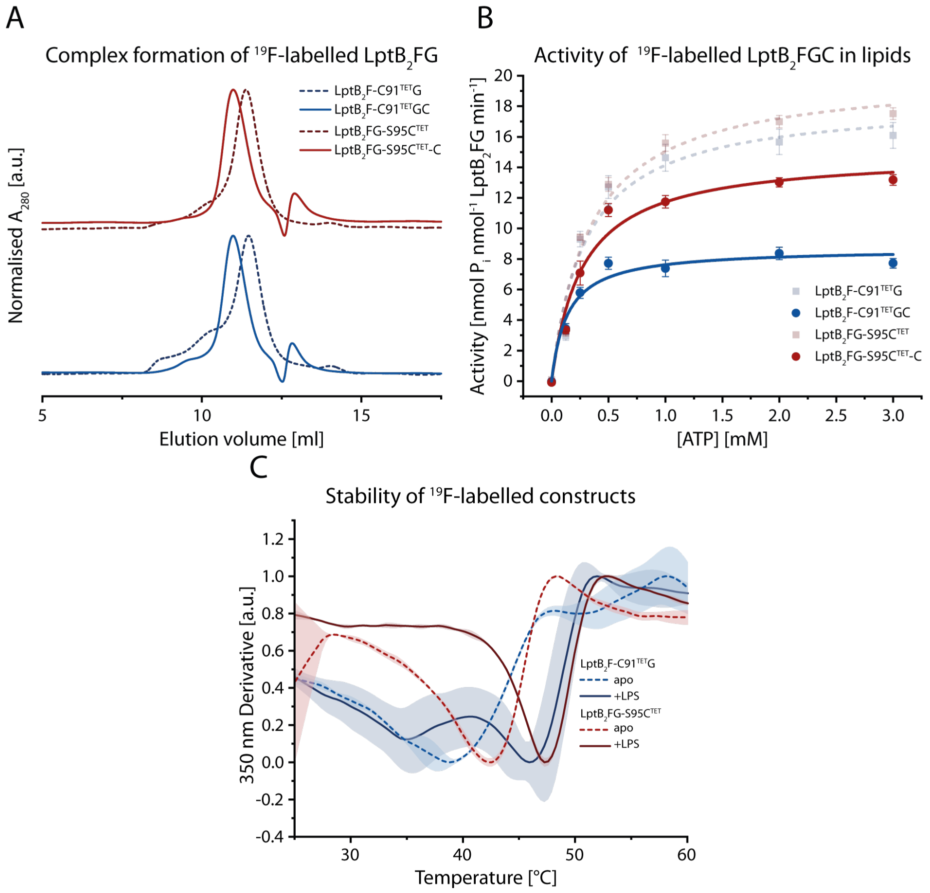


**Figure S1:** Biochemical characterization of labelled $LptB_2FG$(C) constructs**. (A)** Size exclusion chromatography of CH-LptF (blue dashed line) and CH-LptG (red dashed line) and re-solubilized CH-LptFC (blue solid line) and CH-LptGC (red solid line). **(B)** ATPase assay of TET-labelled CH-LptF (blue dashed line), CH-LptG (red dashed line) and in complex with LptC (blue and red solid line) in POPE/POPG liposomes. **(C)** NanoDSF measurement of labelled $LptB_2FG$ constructs. The first derivative of the fluorescence emission at 350 nm is shown in dependence of temperature. CH-LptF (blue) and CH-LptG (red) were measured in detergent micelles in absence (dashed lines) and presence of tenfold molar excess of LPS (solid lines).

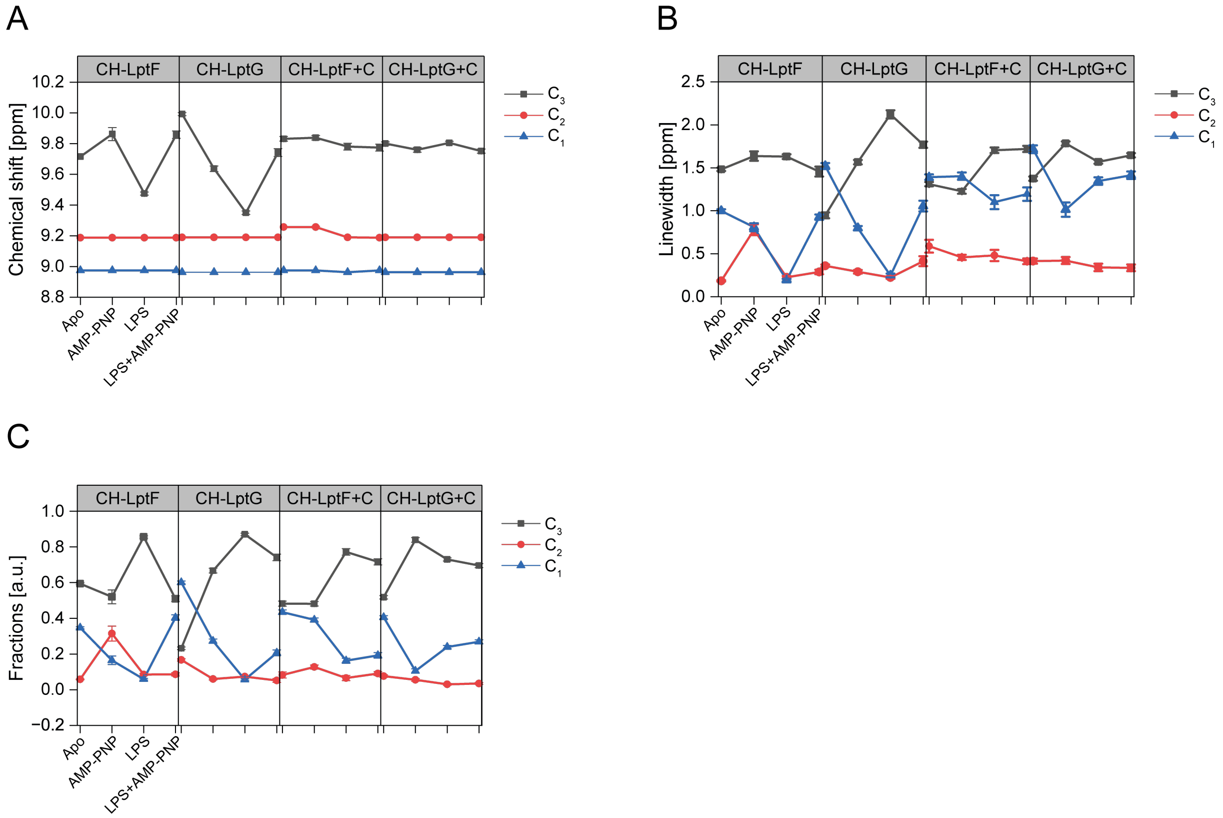


**Figure S2:** Progression of the deconvolution parameters for the C1, C2, and C3 peaks across all experimentally prepared protein states (see Figures 2 and 3 and Tables S1–S4). **(A)** Chemical shift changes. **(B)** Spectral linewidths. **(C)** Relative peak intensities.

**Table S1:** Deconvolution results for the spectra of $LptB_2F(C91^{TET})G$ (CH-LptF) (Fig. 2). The spectral lineshape was fitted with three Lorentzian peaks C1, C2 and C3. The fraction value describes the relative proportion of each Lorentzian peak integral to integral sum of all peaks. The errors are obtained from the peak-fitting routine (see Materials and Methods).

| | C3 | | | | C2 | | | | C1 | | | |
|---|---|---|---|---|---|---|---|---|---|---|---|---|
| | $\delta(^{19}F)$ [ppm] | $\nu_{1/2}$ [ppm] | Integral [a.u.] | Fraction [a.u.] | $\delta(^{19}F)$ [ppm] | $\nu_{1/2}$ [ppm] | Integral [a.u.] | Fraction [a.u.] | $\delta(^{19}F)$ [ppm] | $\nu_{1/2}$ [ppm] | Integral [a.u.] | Fraction [a.u.] |
| **Apo** | 9.72 ± 0.01 | 1.48 ± 0.02 | 1.05 ± 0.01 | 0.59 ± 0.01 | 9.19 | 0.18 ± 0.01 | 0.1 ± 0.01 | 0.06 ± 0.01 | 8.97 ± 0.01 | 1 ± 0.02 | 0.61 ± 0.01 | 0.35 ± 0.01 |
| **AMP-PNP** | 9.86 ± 0.04 | 1.64 ± 0.05 | 1.02 ± 0.09 | 0.52 ± 0.04 | 9.19 | 0.78 ± 0.06 | 0.62 ± 0.11 | 0.31 ± 0.05 | 8.97 | 0.81 ± 0.05 | 0.32 ± 0.05 | 0.16 ± 0.03 |
| **LPS** | 9.47 ± 0.01 | 1.63 ± 0.03 | 1.43 ± 0.02 | 0.86 ± 0.01 | 9.19 | 0.23 ± 0.02 | 0.14 ± 0.01 | 0.09 ± 0.01 | 8.97 | 0.19 ± 0.01 | 0.1 ± 0.01 | 0.06 ± 0.01 |
| **LPS & AMP-PNP** | 9.86 ± 0.02 | 1.46 ± 0.06 | 0.87 ± 0.05 | 0.51 ± 0.02 | 9.19 | 0.29 ± 0.03 | 0.15 ± 0.02 | 0.09 ± 0.01 | 8.97 | 0.93 ± 0.03 | 0.69 ± 0.03 | 0.4 ± 0.02 |

**Table S2:** Deconvolution results for the spectra of $LptB_2FG(S95C^{TET})$ (CH-LptG) (Fig. 2). The spectral lineshape was fitted with three Lorentzian peaks C1, C2 and C3. The fraction value describes the relative proportion of each Lorentzian peak integral to integral sum of all peaks.

| | C3 | | | | C2 | | | | C1 | | | |
|---|---|---|---|---|---|---|---|---|---|---|---|---|
| | $\delta(^{19}F)$ [ppm] | $\nu_{1/2}$ [ppm] | Integral [a.u.] | Fraction [a.u.] | $\delta(^{19}F)$ [ppm] | $\nu_{1/2}$ [ppm] | Integral [a.u.] | Fraction [a.u.] | $\delta(^{19}F)$ [ppm] | $\nu_{1/2}$ [ppm] | Integral [a.u.] | Fraction [a.u.] |
| **Apo** | 9.99 ± 0.01 | 0.95 ± 0.03 | 0.37 ± 0.01 | 0.23 ± 0.01 | 9.19 | 0.36 ± 0.01 | 0.27 ± 0.01 | 0.17 ± 0.01 | 8.96 | 1.52 ± 0.03 | 0.97 ± 0.02 | 0.6 ± 0.01 |
| **AMP-PNP** | 9.64 ± 0.02 | 1.57 ± 0.03 | 1.28 ± 0.05 | 0.67 ± 0.02 | 9.19 | 0.29 ± 0.02 | 0.12 ± 0.01 | 0.06 ± 0.01 | 8.96 | 0.8 ± 0.03 | 0.52 ± 0.03 | 0.27 ± 0.02 |
| **LPS** | 9.35 ± 0.01 | 2.12 ± 0.05 | 1.74 ± 0.03 | 0.87 ± 0.01 | 9.19 | 0.22 ± 0.02 | 0.15 ± 0.01 | 0.07 ± 0.01 | 8.96 | 0.25 ± 0.02 | 0.11 ± 0.01 | 0.06 ± 0.01 |
| **LPS & AMP-PNP** | 9.74 ± 0.03 | 1.77 ± 0.04 | 1.92 ± 0.09 | 0.74 ± 0.02 | 9.19 | 0.41 ± 0.06 | 0.13 ± 0.03 | 0.05 ± 0.02 | 8.96 | 1.05 ± 0.06 | 0.54 ± 0.05 | 0.21 ± 0.02 |

**Table S3:** Deconvolution results for the spectra of LptB$_2$F(C91$^{TET}$)GC (CH-LptF+C) (Fig. 3). The spectral lineshape was fitted with three Lorentzian peaks C1, C2 and C3. The fraction value describes the relative proportion of each Lorentzian peak integral to integral sum of all peaks.

| | C3 | | | | C2 | | | | C1 | | | |
|---|---|---|---|---|---|---|---|---|---|---|---|---|
| | δ($^{19}$F) [ppm] | $\nu_{1/2}$ [ppm] | Integral [a.u.] | Fraction [a.u.] | δ($^{19}$F) [ppm] | $\nu_{1/2}$ [ppm] | Integral [a.u.] | Fraction [a.u.] | δ($^{19}$F) [ppm] | $\nu_{1/2}$ [ppm] | Integral [a.u.] | Fraction [a.u.] |
| **Apo** | 9.83 ± 0.02 | 1.31 ± 0.02 | 1.19 ± 0.05 | 0.48 ± 0.02 | 9.26 | 0.59 ± 0.07 | 0.2 ± 0.05 | 0.08 ± 0.02 | 8.97 | 1.39 ± 0.03 | 1.07 ± 0.02 | 0.43 ± 0.02 |
| **AMP-PNP** | 9.84 ± 0.01 | 1.23 ± 0.03 | 0.98 ± 0.04 | 0.48 ± 0.02 | 9.26 ± 0.01 | 0.46 ± 0.03 | 0.26 ± 0.03 | 0.13 ± 0.02 | 8.97 | 1.4 ± 0.04 | 0.8 ± 0.02 | 0.39 ± 0.02 |
| **LPS** | 9.78 ± 0.02 | 1.71 ± 0.03 | 2.05 ± 0.09 | 0.77 ± 0.02 | 9.19 | 0.48 ± 0.07 | 0.17 ± 0.04 | 0.07 ± 0.02 | 8.96 | 1.1 ± 0.08 | 0.43 ± 0.04 | 0.16 ± 0.02 |
| **LPS & AMP-PNP** | 9.77 ± 0.02 | 1.72 ± 0.04 | 1.72 ± 0.08 | 0.72 ± 0.02 | 9.19 | 0.41 ± 0.03 | 0.22 ± 0.03 | 0.09 ± 0.02 | 8.97 | 1.19 ± 0.08 | 0.46 ± 0.04 | 0.19 ± 0.02 |

**Table S4:** Deconvolution results for the spectra of LptB2FG(S95C$^{TET}$)C (CH-LptG+C) (Fig. 3). The spectral lineshape was fitted with three Lorentzian peaks C1, C2 and C3. The fraction value describes the relative proportion of each Lorentzian peak integral to integral sum of all peaks.

| | C3 | | | | C2 | | | | C1 | | | |
|---|---|---|---|---|---|---|---|---|---|---|---|---|
| | δ($^{19}$F) [ppm] | $\nu_{1/2}$ [ppm] | Integral [a.u.] | Fraction [a.u.] | δ($^{19}$F) [ppm] | $\nu_{1/2}$ [ppm] | Integral [a.u.] | Fraction [a.u.] | δ($^{19}$F) [ppm] | $\nu_{1/2}$ [ppm] | Integral [a.u.] | Fraction [a.u.] |
| **Apo** | 9.8 ± 0.01 | 1.37 ± 0.02 | 1.26 ± 0.04 | 0.52 ± 0.01 | 9.19 | 0.42 ± 0.03 | 0.19 ± 0.02 | 0.08 ± 0.01 | 8.96 | 1.72 ± 0.04 | 0.99 ± 0.02 | 0.41 ± 0.01 |
| **AMP-PNP** | 9.76 ± 0.02 | 1.78 ± 0.02 | 2.3 ± 0.07 | 0.84 ± 0.02 | 9.19 | 0.42 ± 0.04 | 0.15 ± 0.02 | 0.06 ± 0.01 | 8.96 | 1.01 ± 0.08 | 0.29 ± 0.03 | 0.11 ± 0.02 |
| **LPS** | 9.81 ± 0.01 | 1.57 ± 0.02 | 2.02 ± 0.04 | 0.73 ± 0.01 | 9.19 | 0.34 ± 0.04 | 0.09 ± 0.01 | 0.03 ± 0.01 | 8.96 | 1.35 ± 0.04 | 0.66 ± 0.03 | 0.24 ± 0.01 |
| **LPS & AMP-PNP** | 9.75 ± 0.01 | 1.65 ± 0.02 | 1.87 ± 0.05 | 0.7 ± 0.02 | 9.19 | 0.34 ± 0.04 | 0.09 ± 0.01 | 0.04 ± 0.01 | 8.96 | 1.41 ± 0.04 | 0.72 ± 0.03 | 0.27 ± 0.02 |
